# Single-cell RNA sequencing reveals molecular features of postnatal maturation in the murine retinal pigment epithelium

**DOI:** 10.1101/2022.05.07.491046

**Authors:** Ravi S. Pandey, Mark P. Krebs, Mohan T. Bolisetty, Jeremy R. Charette, Jürgen K. Naggert, Paul Robson, Patsy M. Nishina, Gregory W. Carter

**Affiliations:** The Jackson Laboratory, 600 Main Street, Bar Harbor ME 04609, USA

**Keywords:** mouse models of eye disease, cluster analysis

## Abstract

Transcriptomic analysis of the mammalian retinal pigment epithelium (RPE) aims to identify cellular networks that influence ocular development, maintenance, function, and disease. However, available evidence points to RPE cell heterogeneity in the native tissue, which adds complexity to transcriptomic analysis. Here, to assess cell heterogeneity, we performed single-cell RNA sequencing of RPE cells from two young adult male C57BL/6J mice. Following quality control to ensure robust transcript identification limited to cell singlets, we detected 13,858 transcripts among 2,667 and 2,846 RPE cells, respectively. Dimensional reduction by principal component analysis and uniform manifold approximation and projection revealed six distinct cell populations. All clusters expressed transcripts typical of RPE cells; the smallest (C1, containing 1–2% of total cells) exhibited hallmarks of stem and/or progenitor cells. Placing C1–6 along a pseudotime axis suggested a relative decrease in melanogenesis and stem/progenitor gene expression, and a corresponding increase in visual cycle gene expression upon RPE maturation. K-means testing of all detected transcripts identified additional expression patterns that may advance understanding of RPE stem/progenitor cell maintenance and the evolution of cellular metabolic networks during development. This work provides new insights into the transcriptome of the mouse RPE and a baseline for identifying experimentally induced transcriptional changes in future studies of this tissue.

## 1. Introduction

Cells of the retinal pigment epithelium (RPE), an epithelial monolayer located between the neurosensory retina and the choriocapillaris, perform activities that are critical to ocular development and visual function [1-3]. As part of the outer blood-retinal barrier (BRB), RPE cells control the flow of electrolytes, water, gases, nutrients, and waste products between the retina and circulation that is essential for retinal development and homeostasis [1-4]. RPE cells phagocytose the tips of photoreceptor outer segments and thereby contribute to the daily turnover of the phototransduction machinery that maintains visual function [5, 6]. These cells contribute directly to vision by regulating the concentration of ions in the subretinal space, which influence light-dependent electrophysiological responses in photoreceptor cells [7]. They also participate in the visual cycle, in which all-*trans* retinaldehyde released by photoreceptor cells upon light stimulation is reisomerized to the 11-*cis* configuration required for detecting additional stimuli [8]. RPE cells are heavily pigmented with melanin, which absorbs light to improve visual contrast and scavenges reactive oxygen species to maintain tissue homeostasis [9-11]. Although many genes and gene products that contribute to these functions are known, further studies are needed for a molecular understanding of how the RPE contributes to vision, retinal homeostasis, posterior eye development, and ocular disease. RPE transcriptome analysis of native tissue represents a primary approach to this end.

The RPE cell population in the mammalian eye is heterogeneous [12], which may add complexity to transcriptomic analysis. Heterogeneity of RPE cell morphological features as observed in the native tissue, such as cell area, shape, melanin pigmentation, and the number of nuclei per cell, is well documented in human [13-22] and other mammalian species [13, 23, 24], including mouse [25, 26]. Depending on species, morphological differences among RPE cells are accentuated with age and vary topographically with respect to ocular region (central-peripheral, dorsal-ventral, nasal-temporal) and proximity to ocular specializations, including the macula, *area centralis*, visual streak, and tapetum lucidum [23]. Additionally, morphological differences have been observed in adjacent RPE cells or small patches of cells independent of topographical location, resulting in cellular mosaicism [12]. RPE cells are also functionally heterogeneous. For example, although the adult RPE is largely post-mitotic, studies of human eyes have identified a small population of stem cells that proliferate and differentiate when subsequently cultured *in vitro* [27]. Similarly, rare cells containing mitotic figures have been identified in the adult albino rat RPE [13], and a small population of mitotically active cells has been reported in the peripheral RPE of the adult rat [28, 29], which may be related to human RPE stem cells. Analysis of another functional readout, indocyanine green dye uptake, has revealed cellular mosaicism in the human and mouse RPE [30, 31]. Further evidence of heterogeneity has come from histological analysis of cellular components in ocular sections or RPE-choroid-sclera flatmounts, or from biochemical analysis of dissected regions of the posterior eye [12]. Finally, focal RPE changes have been observed in individuals affected with inherited eye diseases, such as the macular dystrophies butterfly shaped pigment (or pattern) dystrophy [32], and Best vitelliform macular dystrophy [33], and in animal models of these diseases [32, 34]. The non-uniform distribution of pathological changes in these diseases raises the possibility of a heterogeneous RPE response to the genetic and/or environmental conditions that induce disease. Overall, these studies provide compelling evidence for topographic and cellular heterogeneity of the RPE. However, the underlying mechanisms that lead to this heterogeneity are poorly understood and may benefit from transcriptomic approaches that provide information at the singlecell level.

Transcriptomic studies have been described using RPE preparations from human donors and mouse models (Supplementary Table 1). Microarray analysis, based on hybridization of known genes, has provided initial insights. RNA sequencing (RNAseq) was used subsequently to improve quality, depth of sequencing, and provide a view of absolute transcript abundance. Most recently, single-cell RNA sequencing (scRNA-seq) has been used to examine cellular heterogeneity in human RPE and provide clues to the development of this tissue [35, 36].

Here, we apply scRNA-seq to RPE cells isolated directly from the mouse posterior eyecup by enzymatic and mechanical disaggregation and by further selection for viability based on fluorescence-activated cell sorting. A transcriptome of about 2,000 highly variable genes was documented in each of roughly 2,700 cells, enabling the use of cluster analysis to identify distinct but related RPE cell populations that appear to be distributed along a maturation time course. Bioinformatic analysis identified major known RPE pathways, including the visual cycle and melanogenesis genes, as well as additional transport and metabolic pathways that appear to be coordinately regulated upon RPE maturation. The approach described may benefit future efforts to understand the molecular basis of RPE function in vision, development, homeostasis, and ocular disease.

## 2. Results

### 2.1. Preparation of single RPE cells

To assess the cellular and molecular heterogeneity of RPE cells in an unbiased manner, we performed scRNA-seq on cell samples obtained from the RPE of two young adult male C57BL/6J (B6) mice (replicates R1 and R2) at postnatal day 36 (P36). This age is three weeks past the last major wave of RPE cell division, which completes at about P15 [25]. To obtain single RPE cells, RPE-choroid-sclera eyecups were incubated with a concentrated trypsin solution and agitated gently to release RPE sheets, which were then disrupted mechanically to enrich for single RPE cells (Figure 1A). Cells from both eyes of each mouse analyzed were pooled and incubated with calcein AM to mark viable cells. In the experiment described here, fluorescence-activated cell sorting (FACS) of the R1 and R2 preparations yielded 27,657 and 12,540 single viable cells, respectively. These yields correspond to 26–28% and 12–13%, respectively, of the total population based on estimates of 5.4 × 104 and 4.9 × 104 RPE cells per adult B6 eye [25, 26]. Single cells obtained by this approach were heavily pigmented and often exhibited two lobes of variable size (Figure 1C). This shape is consistent with an apical and basal cellular domain separated by a junctional actin band as observed in other studies, in which single RPE cells were isolated from native or cultured sheets [37-39]. FACS-purified cells were concentrated and applied to the wells of an scRNA-seq chip to create a bar-coded cDNA library for sequencing (Figure 1A). The elapsed time between enucleation and loading RPE cells onto the scRNA-seq chip was roughly 2 h; cells were kept on ice during this period except for the 30 min trypsin incubation at 37 °C and the 15 min calcein-AM incubation at room temperature.

**Figure 1.**
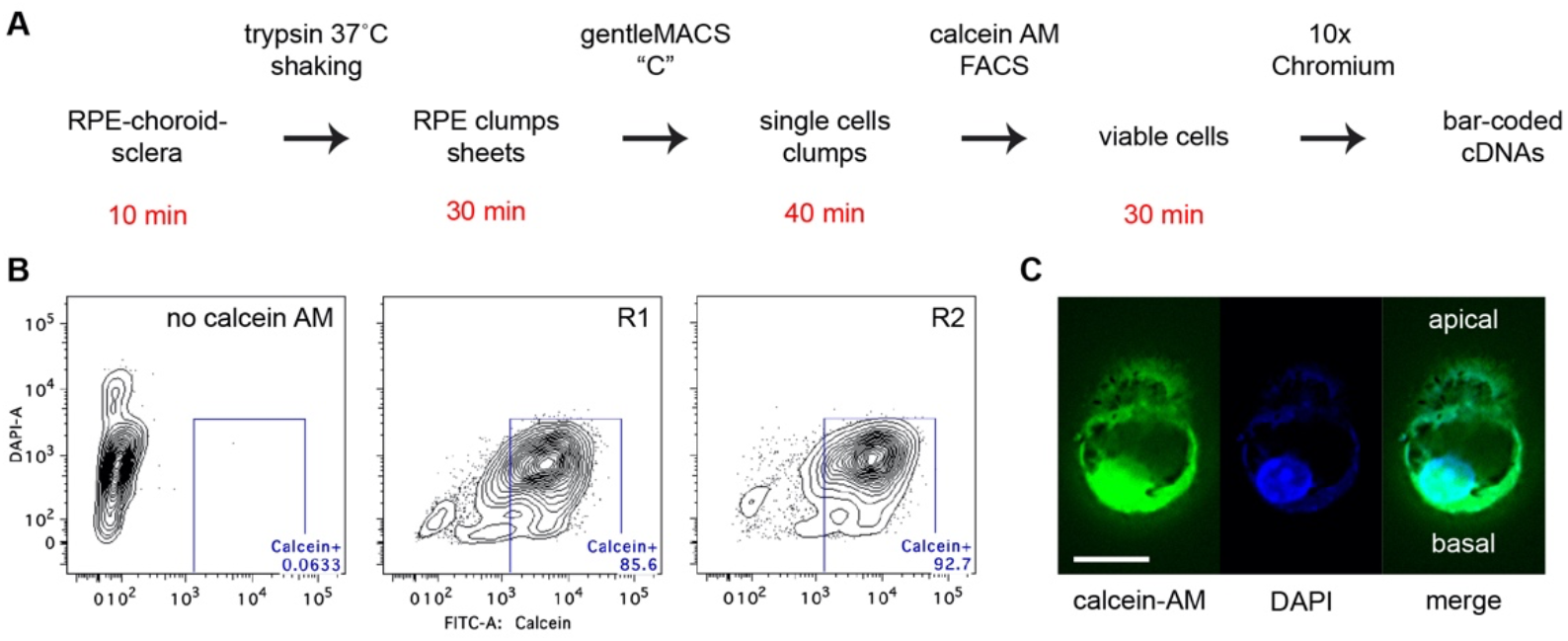
Single RPE cell isolation. (**A**) Flow chart of cell isolation procedure. The time required to complete each step is indicated in red. (**B)** Fluorescence activated cell sorting of RPE preparations labeled with DAPI alone (left panel) or with calcein AM to detect viable cells (middle and right panels). (**C**) Immunofluorescence of viable single cells stained with calcein AM and DAPI. Two lobes are evident, consistent with polarized epithelial cells containing apical and basal domains separated by a junctional actomyosin band. Pigment granules can be identified in both lobes and a nucleus is present in the basal lobe. Scale bar, 10 µm.

### 2.2. Characterization of RPE scRNA-seq datasets

A total of 2667 and 2846 cells from replicates R1 and R2, respectively, that passed all quality checks (see Methods) were analyzed. Unsupervised clustering of individual cell transcriptomes using Louvain community detection revealed six transcriptionally distinct clusters in both samples (Fig. 2 A, B, Table 1). By default, the Seurat software labeled these as clusters 0–5 based on the population size of each cluster (Fig. 2C). We renumbered the clusters produced by Seurat as C1–C6, based in part on the clustering tree map produced from the clustering trees tool [40], which shows how cells move between clusters and how unstable clusters split into smaller clusters as clustering resolution is increased. The clustering tree map for single cell data from R1 revealed that cells in Seurat cluster 5 at a final chosen resolution of 0.6 split from the remaining cells at the first step (clustering resolution = 0.1), suggesting a distinct gene expression profile in these cells compared to other cells. Therefore, we renumbered Seurat cluster 5 as cluster C1 (Fig. 2C). On the other hand, Seurat clusters 1–3 appeared at later steps of this analysis (clustering resolution = 0.3, 0.6), so we renumbered these as clusters C4–C6 (Fig. 2C). Finally, Seurat clusters 0 and 4 were labelled as C2 and C3, respectively. The rationale for final adjustment of the cluster order is given in section 2.4.

**Figure 2.**
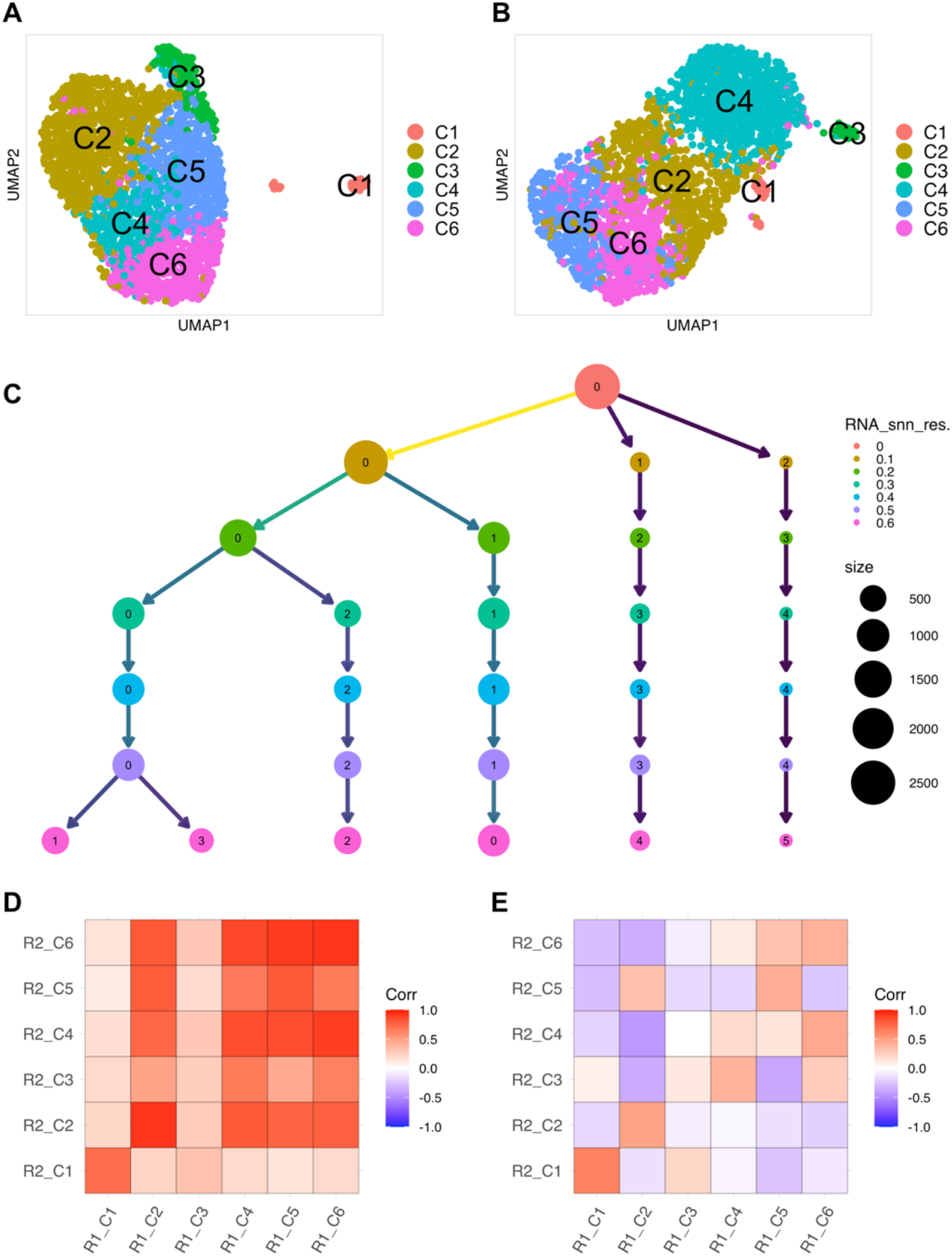
Single-cell transcriptomic analysis of mouse RPE. (**A**) UMAP projection of 2667 single cells obtained from R1 RPE cells. Data are shown in two dimensions using UMAP. Unsupervised analysis clustered cells into six transcriptionally distinct populations, each plotted in a different color. (**B**) UMAP projection of 2846 single cells obtained from R2 RPE cells displayed as in **A**. (**C**) Clustering tree of 2667 single cells from R1. Results from clustering using Seurat with resolution parameters of 0–0.6. At a resolution of 0.1, three main branches are observed, one of which continues to split up to a resolution of 0.6 while the other two remain intact. Seurat labels clusters according to their size, with cluster 0 being the largest. Clusters 5, 0, 4, 3, 2, and 1 were relabeled as C1, C2, C3, C4, C5, and C6, respectively as shown in **A**. (**D**) Pearson correlation between single cell clusters from R1 and R2 using the average expression of genes in each cluster. (**E**) Pearson correlation between single cell clusters from R1 and R2 using log2FC of genes in each cluster relative to the average gene expression in the union of cells from all other clusters. Positive correlations are shown in red and negative correlations in blue. Correlations with nominal p-value < 0.05 were considered significant.

**Table 1.** Cell populations in each cluster identified in R1 and R2.

| Cluster | R1 | R2 |
| --- | --- | --- |
| C1 | 57 | 40 |
| C2 | 931 | 687 |
| C3 | 186 | 52 |
| C4 | 378 | 980 |
| C5 | 556 | 498 |
| C6 | 559 | 589 |

We performed correlation analysis to assess whether the clusters identified in each replicate were distinct and to test whether the replicates were similar. In each replicate, correlation analysis between clusters using average gene expression as a metric indicated strong positive correlation between Clusters C2–6, but weak correlation of these clusters with C1 (Supplementary Figure 1A, C). However, correlation analysis using the logarithm (base 2) of the fold-change in gene expression (log2FC) as a relative expression metric revealed negative correlations among the clusters (Supplementary Figure 1B, D), suggesting that cluster analysis identified distinct but related RPE cell populations. To compare clustering in R1 and R2, we measured the correlation between the average expression of genes in each cluster from both replicates (Figure 2D) as well as the correlation between log2FC in each cluster from both replicates (Figure 2E). We noticed substantial similarity between clusters in R1 and R2. C1 from R1 and R2 correlated strongly with each other, and clusters C2–6 correlated with each other (Figure 2D–E). Taken together, these results suggest that the gene expression profiles in R1 and R2 are generally similar and confirm that the RPE cells isolated by our approach represent a robust heterogeneous population. Results from R1 are presented below, and those for R2 are summarized at the end of the Results section.

### 2.3. Functional Analysis of Clusters

We found ≥ 25 cluster-specific marker genes for each cluster in R1 (adjusted p value [padj] < 0.05, Supplementary File 1). The top 20 marker genes that distinguished C1 from clusters C2–C6 are shown in Figure 3A. C2–C6 were relatively less distinguished from each other, suggesting the RPE cell population is closely related in these clusters. C1 showed a higher expression of genes identified implicated in stem/progenitor maintenance and/or stemness, such as *Aldoc* [41], *Dkk3* [42, 43], and *Id3* [41, 44, 45], compared to C2–C6. By contrast, C2–C6 exhibited increased expression of RPE-specific marker genes such as the visual cycle genes *Rpe65, Lrat*, and *Rrh* [8], compared to C1 (Figure 3A). Further, gene ontology (GO) analysis identified enrichment of melanin biosynthetic and energy metabolism related biological processes in differentially upregulated genes (padj < 0.05) in C1, while differentially upregulated genes (padj < 0.05) in C2–C6 were enriched for multiple biological processes as indicated by GO terms related to Wnt signaling, cellular response to metal ions, detoxification, and others (Figure 3B; Supplementary File 2).

**Figure 3.**
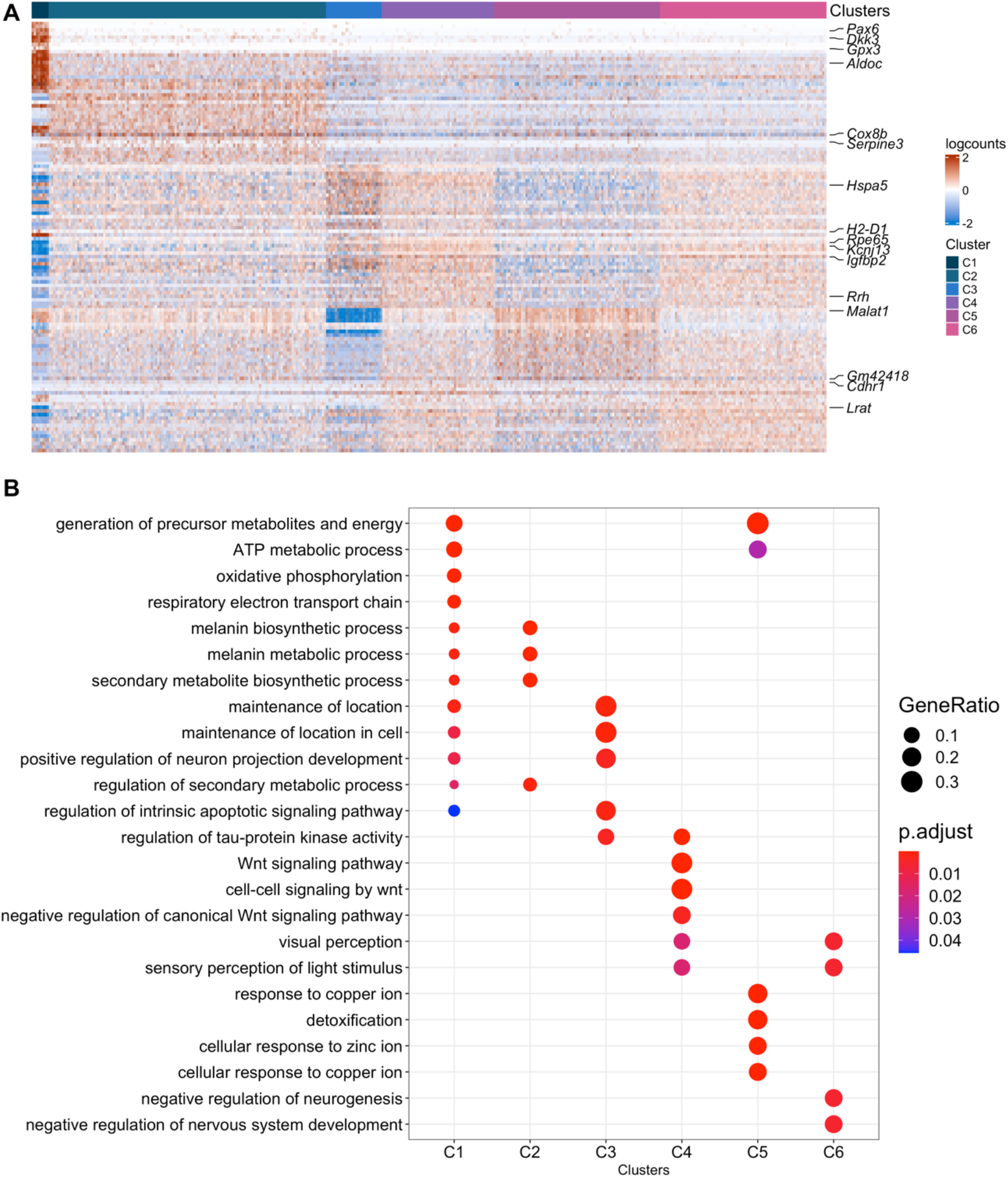
Differential Expression Analysis of RPE cell clusters in R1. (**A**) The top 20 differentially expressed genes in clusters, ranked by false discovery rate (FDR), are shown in the heatmap. Gene expression values were centered, scaled, and transformed to a scale from −2 to 2. Select signature genes are highlighted on the right. (**B**) Enrichment of biological processes in differentially upregulated genes in each cluster using clusterprofiler. The significance threshold for all enrichment analyses was set to 0.05 using Benjamini-Hochberg corrected p-values.

### 2.4. Assessment of Possible Retinal or Choroidal Cell Contamination

To examine the possibility that the cellular heterogeneity present in R1 and R2 was due to contamination by non-RPE cells, we first compared our cell clusters with mouse retinal cell clusters from a previous study [46]. C1 exhibited a significant positive correlation (p < 0.05) with most of the mouse retinal cell clusters, while clusters C2 and C4 did not correlate significantly with mouse retinal cell clusters (Figure 4A). Clusters C5 and C6 exhibited significant negative correlation (p < 0.05) with mouse retinal cell clusters (Figure 4A). The significant positive correlation of cluster C1 with mouse retinal cell clusters from 2-week-old mice [46] suggests that cells in C1 either are not RPE cells or possibly represent a multipotent RPE cell type that expresses ret-ina-associated genes.

**Figure 4.**
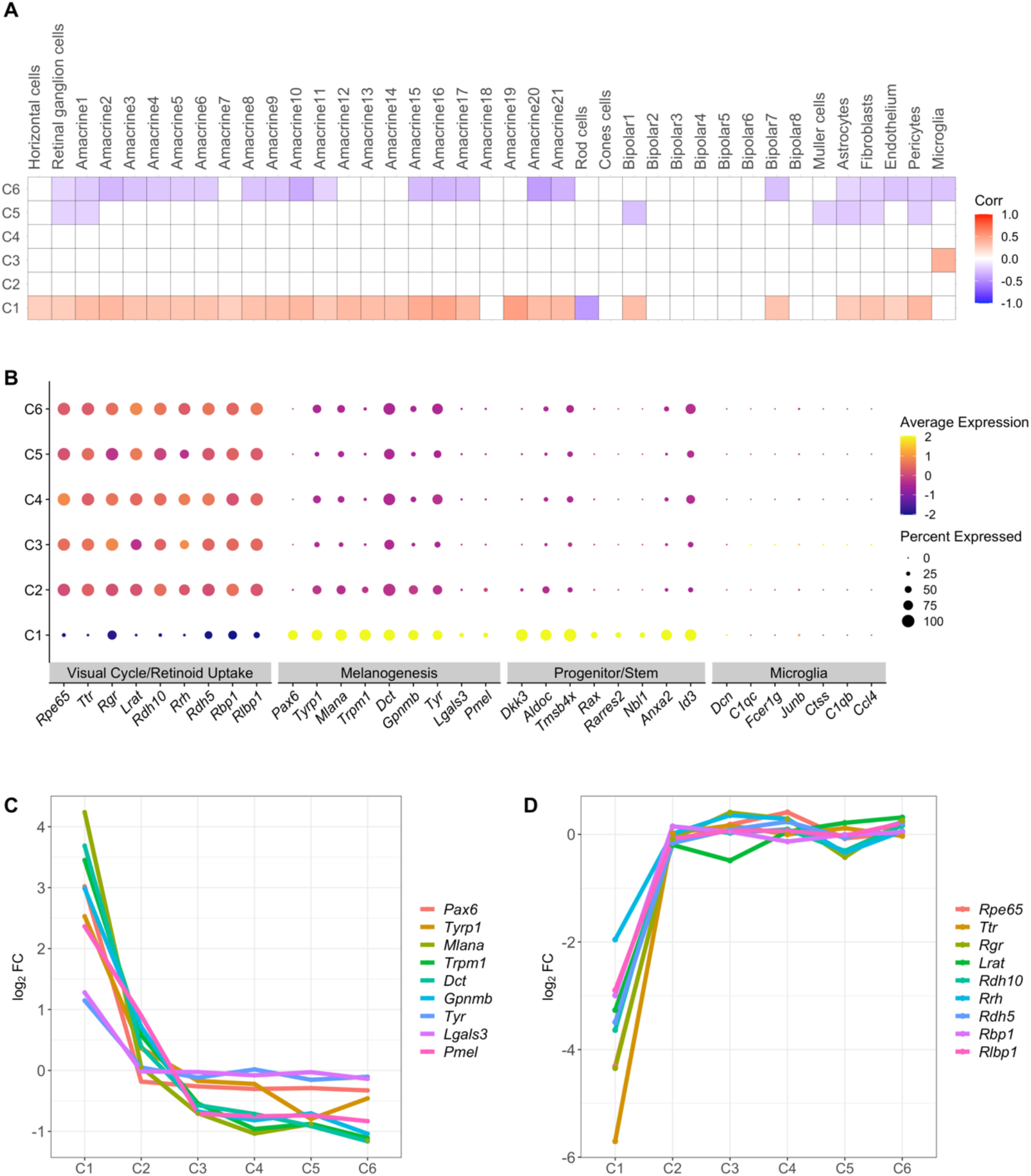
Heterogeneity of RPE cell populations from R1. (**A**) Correlation between single cell clusters in R1 and microglial retinal cell clusters. Pearson correlation coefficients were calculated for log2FC of genes in each cluster. Positive correlations are shown in red and negative correlations in blue. Correlation with nominal p-value < 0.05 are considered significant and shown in figure. (**B**) Dot plot showing marker gene expression for different RPE specific pathways (visual cycle, melanogenesis), and cell types (stem/progenitor cell and immune cells). Dot sizes indicate the percentage of cells in each cluster expressing the gene, and colors indicate average expression levels. (**C**)Differential expression (log2FC) of melanogenesis genes along RPE clusters C1–6 (**D**)Differential expression (log2FC) of visual cycle genes along RPE clusters C1–6.

To distinguish between these possibilities, we examined the expression of genes associated with known RPE pathways such as visual cycle and melanogenesis. Human RPE stem cells have been associated with low levels of visual cycle genes, high levels of melanin pigment biosynthetic genes and stem/progenitor/stem cell markers [27]. Similarly, low expression of visual cycle genes and high levels of melanogenesis and stem/progenitor cell marker genes were observed in C1 compared to C2–6 (Figure 4B). These gene expression results support the assignment of C1 as a stem/progenitor cell type and provide further evidence for heterogeneity of the RPE cell population.

As an additional test of whether C1 and C2–6 represent *bona fide* RPE cells, rather than possible contaminating cell types, we investigated the expression of other genes in the cell clusters. Microglia are sometimes observed at the interface between the retina and RPE of B6 mice [47], and therefore are a possible contaminating cell type. We identified very low levels of key microglial cell type marker genes in all clusters (Figure 4B). We also implemented the CELL-ID method [48] to verify the identity of the cell clusters using marker genes of microglial cell type as well as marker genes reported for the microglial cluster in the mouse retina [46] as a reference. CELL-ID annotated only five cells as microglia in R1 (Table 2), indicating that there was little contamination from these cells in this dataset (<0.2% of total, <3% of any cluster). In R2, 26 microglia were identified, but remain a small percentage of the cell population (<1% of total, <3% of any cluster). Next, we examined the possible presence of choroidal melanocytes cells, as melanogenesis related genes, such as *Pmel* and *Mlana*, which were highly expressed from cell cluster C1 (Figure 4B; Supplementary Figure 2C), are also expressed in both mouse and human choroidal melanocytes [49, 50]. We compared the top 100 gene signatures of each cell in C1–C6 with those in choroidal melanocytes from a mouse study [49] using the CELL-ID approach [48]. This approach identified eight cells in C2 with a significant choroidal melanocyte gene signature in R1, and one cell in C2 of R2 (Table 5; <0.4% of total, <1% of any cluster). None of the cells in cluster C1 were identified with this signature. Taken together, these results suggest that most of the cells in all clusters, including C1, correspond to RPE cells and that contamination with microglia or choroidal melanocytes is rare.

**Table 2.** Number of cells in each RPE cluster identified as melanocyte and microglial 265 cell types using CELL-ID.

| Cluster | R1 |  | R2 |  |
| --- | --- | --- | --- | --- |
|  | melanocytes | microglia | melanocytes | microglia |
| C1 | 0 | 0 | 0 | 1 |
| C2 | 8 | 0 | 1 | 0 |
| C3 | 0 | 4 | 0 | 0 |
| C4 | 0 | 0 | 0 | 23 |
| C5 | 0 | 0 | 0 | 0 |
| C6 | 0 | 1 | 0 | 2 |

### 2.5. A Proposed Cluster Maturation Timeline

Based on the expression of genes related to RPE-specific pathways such as visual cycle and melanogenesis and correlation with mouse retinal cell clusters (Figure 4A, B), we propose C1 consists of immature RPE cells and C2–6 contain mature RPE cell types. We staged a possible maturation timeline from C1 to C6 based on well-known attributes of RPE cells (Figure 4C, D). Melanogenesis-associated genes exhibited significantly increased expression (log2FC > 1; padj < 0.05) in C1 relative to other clusters (Table 3, Figure 4C) and their expression declined progressively with maturation from C2–6 (Figure 4C). Similarly, the expression of visual cycle and retinoid uptake genes was significantly reduced (log2FC < -1; padj < 0.05) in cluster C1 relative to other clusters (Table 4, Figure 4 D). We also examined genes known to contribute to stem/progenitor cell maintenance and renewal or are differentially upregulated in stem/progenitor cells, including *Aldoc, Dkk3, Id3, Tmsb4x* [51], *Anxa2* [52], *Nbl1* [53], *Rax* [54], and *Rarres2* [55]. Transcripts from these genes were more abundant in cluster C1 (Figure 4B, Table 5) and declined in C2–6. Overall, these results support the identification of C1 as a stem/progenitor cell population and the proposed maturation of RPE cells from C1 to C6.

**Table 3.** Differential expression of selected melanogenesis genes in cluster C1.

| Symbol | pathway | R1 |  | R2 |  |
| --- | --- | --- | --- | --- | --- |
| | | $\log_2FC$ | $p_{adj}$ | $\log_2FC$ | $p_{adj}$ |
| *Mlana* | melanogenesis | 4.24 | 2.28E-25 | 3.27 | 1.64E-02 |
| *Dct* | melanogenesis | 3.68 | 2.55E-18 | 2.89 | 1.51E-04 |
| *Trpm1* | melanogenesis | 3.45 | 3.36E-41 | 2.09 | 8.60E-12 |
| *Gpnmb* | melanogenesis | 2.98 | 1.83E-17 | 2.50 | 1.35E-01 |
| *Tyrp1* | melanogenesis | 2.53 | 1.22E-19 | 2.01 | 7.54E-02 |
| *Pmel* | melanogenesis | 2.36 | 2.87E-02 | 2.45 | 4.07E-01 |
| *Lgals3* | melanogenesis | 1.28 | 2.75E-05 | 1.95 | 1.53E-07 |
| *Tyr* | melanogenesis | 1.14 | 3.25E-03 | 2.01 | 7.54E-02 |

**Table 4.**
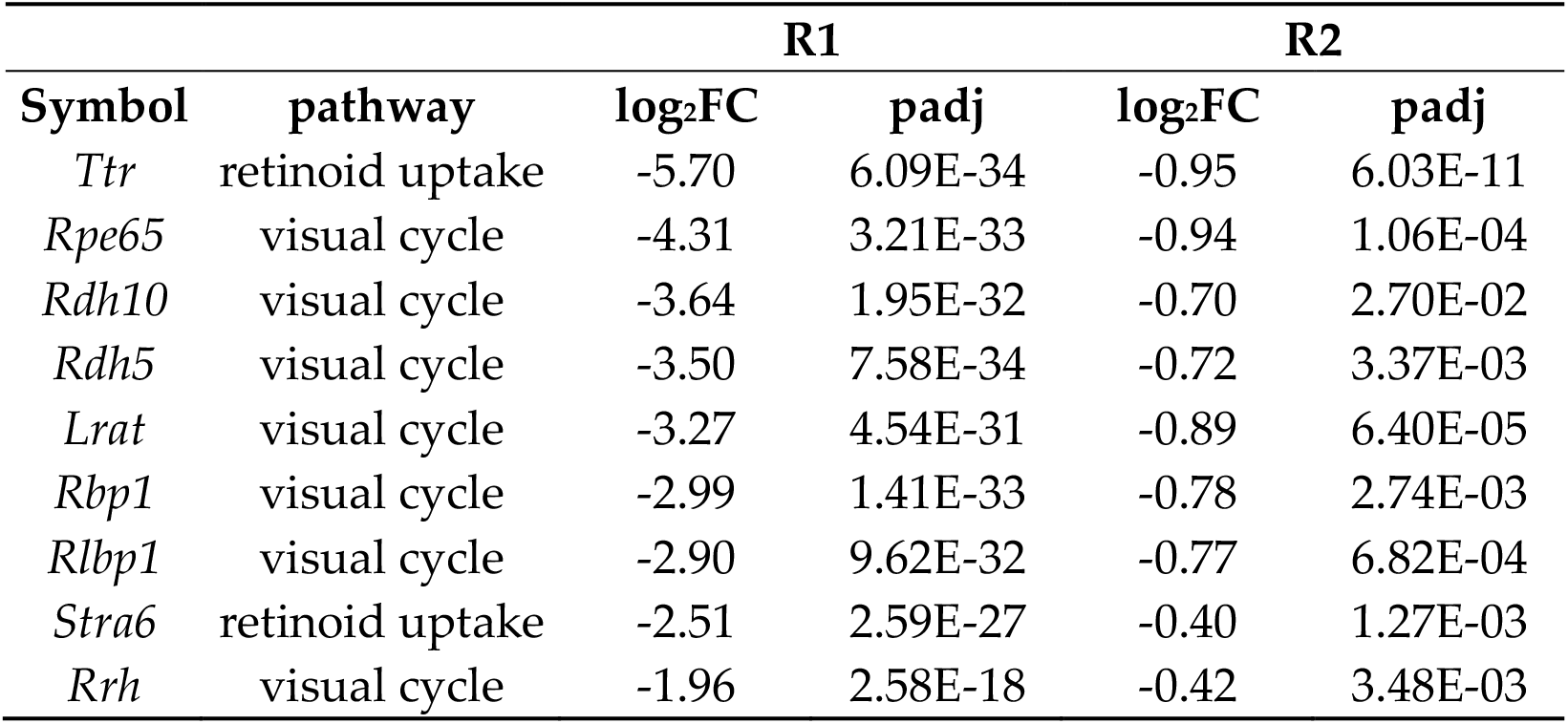
Differential expression of selected visual cycle and retinoid uptake genes in cluster C1.

**Table 5.** Differential expression of selected stem/progenitor cell genes in cluster C1.

| Symbol | pathway | R1 |  | R2 |  |
| --- | --- | --- | --- | --- | --- |
|  |  | log<sub>2</sub>FC | padj | log<sub>2</sub>FC | padj |
| *Aldoc* | maintenance | 4.46 | 1.37E-33 | 3.21 | 7.68E-11 |
| *Tmsb4x* | stemness | 4.09 | 4.57E-41 | 3.52 | 4.27E-06 |
| *Dkk3* | stemness | 3.84 | 2.49E-104 | 3.05 | 3.08E-46 |
| *Id3* | stemness | 3.13 | 2.42E-26 | 2.78 | 1.16E-02 |
| *Anxa2* | maintenance | 2.82 | 5.51E-40 | 1.77 | 2.04E-01 |
| *Nbl1* | maintenance | 2.36 | 8.96E-161 | 1.29 | 5.94E-99 |
| *Rax* | maintenance | 2.24 | 1.41E-64 | 1.93 | 1.96E-66 |
| *Rarres2* | stemness | 1.92 | 9.84E-91 | 1.87 | 5.68E-37 |

### 2.6. k-means Clustering and Functional Profiling

*To identify other biological pathways that exhibit similar trends across the maturation timeline as the melanogenesis, visual cycle, and retinoid uptake pathways, we performed k-means clustering on differentially expressed genes (padj < 0*.*05) across all RPE* clusters. Differentially expressed genes were classified into 10 different groups (Gp1–Gp10) based on their gene expression profiles along the proposed maturation timeline (Supplementary File 3). Of these, gene sets in Gp5 and to a lesser extent Gp2 exhibited an almost identical expression profile as in our initial analysis of visual cycle and retinoid uptake genes along the maturation timeline (that is, downregulated in progenitor C1). In contrast, genes in Gp4 and Gp10 exhibited similar expression profile as melanogenesis genes (that is, upregulated in progenitor C1) along the maturation timeline (Figure 5A). We then performed GO analysis on these gene sets to identify significant enrichment (FDR < 0.05) of multiple biological processes. Genes in Gp2 were enriched for biological processes such as: “regulation of lipid localization”, “transport”, “tissue migration” and “transforming growth factor” (Figure 5B, Supplementary File 4). Genes in Gp5 were enriched for biological processes such as “morphogenesis of an epithelial fold”, “lipid localization”, “homeostasis” and “ERK1 and ERK2 cascade” (Figure 5B, Supplementary File 4). Gene sets in Gp4 were enriched for “metabolic process”, “pigmentation, “establishment of cell polarity”, and “neuron differentiation”. Gp10 genes were enriched for biological processes, such as “oxidative phosphorylation”, “negative regulation of immune system process”, and “epithelial cell proliferation” (Figure 5B, Supplementary File 4). Overall, we identified multiple biological processes that were differentially regulated in proposed progenitor/stem cluster C1.

**Figure 5.**
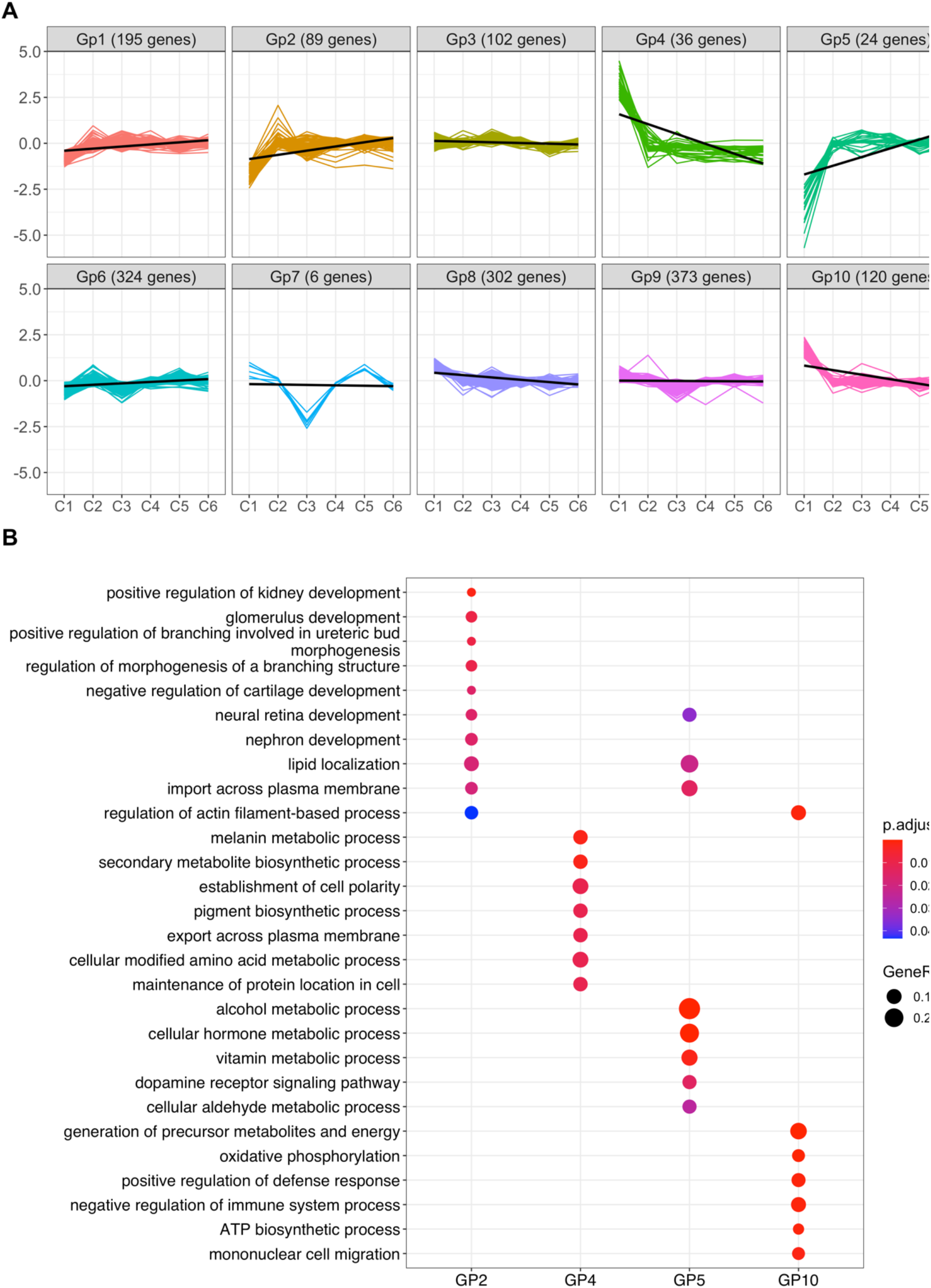
Characterization of RPE subpopulations from R1. (**A**) k-means clustering of differentially expressed genes in RPE cell populations from R1. Clustering analysis identified 10 groups of genes with distinct expression profiles across possible maturation timeline, each plotted in a different color. Number of genes in each group are shown in parentheses (**B**) Enrichment of biological processes in selected groups using clusterprofiler. The significance threshold for all enrichment analyses was set to 0.05 using Benjamini-Hochberg corrected p-values.

### 2.7. Replication of Results in R2

To assess the reproducibility of these findings, we examined results from R2. As described above, unsupervised clustering revealed six transcriptionally distinct clusters in R2, similar to the clustering results in R1 (Figure 2B). Average gene expression was highly correlated in respective clusters from R1 and R2 (Figure 2C). Heatmaps of the top 20 marker genes distinguished C1 from C2–6 (Supplementary Figure 2A) and indicated that C2–6 were relatively less distinguished from each other, suggesting heterogeneous but related RPE cell populations in R2. C1 showed higher expression of stem/progenitor marker genes such as *Dkk3, Id3*, and *Aldoc*. Correlation analysis with mouse retinal cell clusters [46] identified significant positive correlation (p < 0.05) between C1 to most of the mouse retinal cell clusters (Supplementary Figure 2B), while C2 and C4 showed significant negative correlation (p < 0.05) with some of the mouse retinal cell clusters (Supplementary Figure 2B).

Moreover, in R2, C1 expressed high levels of melanogenesis and stem/progenitor cell marker genes and low levels of visual cycle genes, while other clusters exhibited higher levels of visual cycle genes and reduced levels of progenitor markers (Supplementary Figure 2C). A maturation timeline along C1–6 was noted for R2, primarily based on expression profile of melanogenesis genes (Supplementary Figure 2D). However, the expression profile of visual cycle genes was not in complete agreement with this proposed maturation timeline (Supplementary Figure 2E), possibly indicating a lower sample quality in R2. Nonetheless, an overall maturation timeline representing C1 as an immature/progenitor RPE cell type and C2–6 as mature RPE cells was obtained. Importantly, C1 in R2 was highly correlated with C1 in R1 (Figure 2D, E). Thus, we were able to identify stem/progenitor cells in R2 as in R1.

Finally, we also performed k-means clustering on differentially expressed genes in each cluster in R2 followed by GO analysis on group of genes with similar expression profile as melanogenesis and visual cycle genes (Supplementary Figure 3). Groups of genes with reduced expression in C1 but increased levels in mature RPE cells (Gp1 and Gp2) were enriched for “regulation of lipid localization and transport”, “response to toxic substances”, and “transforming growth factor beta production” biological processes (Supplementary Figure 3B, Supplementary File 4). Group of genes with increased expression in C1 but reduced in mature RPE cells (Gp5 and Gp8) were significantly enriched for “metabolic process” and “regulation of transport activity” and “establishment of cell polarity” (Supplementary Figure 3B, Supplementary File 4). Overall, we observed similar cell cluster profiles in R2 and R1, reinforcing the presence of a stem/progenitor population and confirming evidence for RPE cell heterogeneity due to differences in RPE maturation in young adult mice.

### 2.8. A Mature RPE Transcriptome

As indicated above, cluster analysis and k-means testing support the identification of C2–6 as a mature RPE cell population. Importantly, this population is free of contaminating cell types, and its transcriptome may therefore provide an opportunity to characterize the RPE transcriptome with high confidence. To allow such analysis, we determined the average transcript count for each gene in the combined C2–6 “mature” population that passed quality control criteria (Supplementary file 5). Counts from C1 and from individual clusters C2–6 were provided in parallel. A useful (though arbitrary) inclusion threshold is an average expression value of 1 transcript per cell. As expected, signature RPE genes are expressed in the mature population at high levels above this threshold (*Rgr*, 186.9; *Rpe65*, 47.8), whereas photoreceptor genes are expressed below the threshold (*Gnat1*, 0.008; *Rho*, 0.02). Surprisingly, several genes associated with photoreceptors are detected at levels above the threshold (*Rom1*, 4.2; *Slc24a1*, 3.7; *Abca4*, 1.9; *Gnb1*, 1.9), raising the possibility that they are also expressed in the RPE (see Discussion). These data can be queried to assess the presence and abundance of specific gene transcripts levels in the mature RPE cell population.

## 3. Discussion

Transcriptional profiling of the mammalian RPE has been pursued to provide insights into RPE function in ocular health and disease [56, 57]. In this report, we have demonstrated the use of scRNA-seq to assess the transcriptional profile of individual cells obtained from the native RPE of young adult mice. Our results may aid future efforts to explain RPE morphological and functional heterogeneity at the molecular level.

Our results at P36 reveal a major population (>98% of total) of mature RPE cells with overlapping but distinct transcriptomic signatures (C2–6). The clusters are closely related but retain differences in genes associated with known pathways that reflect RPE development, such as the Wnt signaling pathway [58], the visual cycle [8], and the cellular accumulation of copper and other metal ions, which contribute to melanogenesis and ultimately accumulate in melanosomes [59]. C2–6 may represent different stages in the RPE maturation process, which is not synchronized across the full posterior eye. Alternatively, the RPE at P36 may be fully mature, and the observed heterogeneity in gene expression may arise instead from differences in topographic location or from cellular mosaicism.

Our analysis also identifies a small cluster of RPE cells (C1) with possible stem/progenitor cell properties. Stem cells are characterized by self-renewal (the ability to proliferate in an undifferentiated state) and potency (the capacity to yield diverse differentiated states in response to suitable growth and differentiation stimuli) [60, 61]. Progenitor cells are related to stem cells, but their self-renewal is limited to a small number of cell divisions, and their potency is limited to fewer cell types determined by commitment to a specific differentiation pathway [60, 61]. Our results indicate a high correlation of gene expression between C1 and retinal cell clusters, possibly indicating a capacity to differentiate into multiple retinal cell types. High expression of selected stemness and stem/progenitor cell maintenance genes was also observed among C1 cells. These are likely to be RPE cells, as they express melanogenesis genes at high levels, a known attribute of embryonic RPE [62]. Our analysis excluded another melanin-producing cell type, the choroidal melanocyte, as a major constituent of C1. These results suggest that C1 consists of multipotent RPE stem/progenitor cells, which supports prior evidence for such cells in human and rodent eyes [13, 27-29].

GO analysis of clusters and k-means clustering reinforces the identification of C1 and C2–6 as stem/progenitor and mature RPE cells, respectively, and provides new insights into possible RPE stem/progenitor cell function. For interpreting these data, a relevant concept is the existence of a stem-cell niche [63], which protects stem cells from injury due to the surrounding environment and immune system, and which regulates the participation of stem cells in tissue growth, maintenance, and repair. Gp2 (R1) exhibits an overall trajectory of slightly increasing gene expression from C1 to C2–6. A major class of Gp2 GO terms involve development (kidney, glomerulus, cartilage, neural retina, nephron) or morphogenesis (ureteric bud, branching structure). Most of these annotations include the RPE genes *Bmp4, Nog*, and *Sox9*, which encode the secreted growth factor and extracellular matrix (ECM) protein BMP4; its inhibitor NOG, which produces morphogen gradients based on its distribution relative to BMP proteins [64]; and/or the developmental transcription factor SOX9, which regulates ECM production in diverse cell types [65-67]. Other ECM genes are variably associated with Gp2 GO terms, such as collagen genes *Col4a3, Col8a1, Col8a2*. These results suggest tissue growth and ECM production is an important activity of the mature, differentiated RPE that is downregulated in the RPE stem/progenitor niche. A second major class of Gp2 GO terms involve the transport of nutrients across the plasma membrane, dominated by solute carriers for amino acids, lipids, and energy metabolites. These results indicate an altered transport activity in C1 compared to the mature RPE, as might be expected as cells leave the stem cell niche and encounter a new environment.

Genes in the GO terms associated with Gp4 and Gp5 are predominantly those in the melanogenesis pathway and visual cycle, exhibiting decreased and increased expression, respectively, upon RPE maturation from C1 to C2–6. An additional gene of interest associated with diverse Gp4 GO terms describing cell polarity, proliferation, migration, and mitotic spindle orientation is *Gja1*. This gene encodes gap junction protein GJA1 (connexin 43), which participates in multiple pathways in the stem cell niche [68]. Its decrease in expression as cell mature from C1 to C2–6 is consistent with the departure of RPE cells from the stem cell niche. As in Gp2, several GO terms in Gp5 are associated with solute transport across membranes, consistent with an altered transport activity in the RPE stem/progenitor cell population.

Genes associated with Gp10 GO terms indicate a shift in energy metabolism between C1 and C2–6. Gp10 transcripts from genes encoding glycolytic/gluconeogenic enzymes (*Tpi1, Pkm*) as well as mitochondrial components of the tricarboxylic acid cycle (*Sdhb, Idh2*) and oxidative phosphorylation pathway (*Cox4i1, Atp5a1, Atp5c1, Atp5b, Atp5o*) were relatively more abundant in C1 than in C2–6. Interestingly, increased isocitrate dehydrogenase (IDH2) promotes conversion of a-ketoglutarate to citrate by reductive carboxylation, identified as a major RPE metabolic pathway [69]. Reductive carboxylation influences redox homeostasis, which is central to stem cell self-renewal [70, 71], and contributes to RPE fatty acid synthesis [69], which is considered essential for human pluripotent stem cell survival [72]. The altered expression of energy metabolism genes among C1 cells may reflect differences in the local nutrient and redox status of the RPE stem/progenitor cell niche.

Other interesting genes associated with Gp10 GO annotations include *Id1, Tpm1, Pdlim4*, and *Cd47*, which are common to terms involving the assembly of the actin cytoskeleton. TPM1 (tropomyosin 1) regulates actomyosin contraction in muscle and non-muscle cells [73], PDLIM4 promotes actin stress fiber formation [74], and CD47 regulates actin reorganization during induced cell death [75]. ID1 is a transcription factor that mediates cell stemness through negative regulation of basic helix loop helix transcription factors, thereby promoting self-renewal [76, 77]. This protein also promotes stress fiber formation in prostate epithelial cells upon treatment with transforming growth factor ?1 [78]. These studies, together with our results, raise the intriguing possibility that RPE stemness may arise in part from an increased abundance of factors that regulate the actin cytoskeleton.

The use of scRNA-seq provides greater confidence in assigning transcripts to cell types that have traditionally been difficult to isolate or characterize in pure form. In our study, several transcripts were detected in the mature RPE cell population from genes that are thought to be expressed mainly in photoreceptor cells, including as *Abca4, Gnb1, Rom1*, and *Slc24a1*. These transcripts are unlikely to be due to cell contamination but may arise from phagocytic uptake of outer segments. However, it is also conceivable that some or all these genes are functionally expressed in the RPE. A prominent example is *Abca4*, which encodes a membrane flippase that transports vitamin A retinal-lipid adducts in the photoreceptor outer segment and is clinically associated with a form of Stargardt macular dystrophy [79]. The *Abca4* gene and corresponding protein have recently been shown to be expressed both in photoreceptors and in the RPE [80], which may yield new avenues for understanding ocular vitamin A metabolism and interpreting the effect of *ABCA4* mutations in Stargardt disease. Similar investigative opportunities may await other genes expressed in the RPE that have been predominantly characterized as photoreceptor specific. Interestingly, *ROM1* variants cause an *ABCA4-*like macular dystrophy, raising the possibility that ROM1 dysfunction in the RPE contributes to this phenotype [81].

Limitations of this study include the lengthy RPE cell isolation procedure, which may allow changes in RNA levels that prevent accurate determination of *in vivo* RNA abundance. Also, because replicate samples FACS analysis are stored for different lengths of time prior to loading on the single-cell processing system, RNA quality may be compromised. Future refinements that shorten and/or synchronize the isolation procedure to minimize RNA degradation during the procedure may benefit the approach. In addition, the recovery of a relatively small portion of the total RPE cell population may contribute to bias in assessing RPE heterogeneity. Methods that improve RPE sheet disruption may improve the proportion of the total population collected and thereby obtain a more representative cell population. Future efforts include immunohistochemical or *in situ* RNA hybridization studies to validate assess topographic and cellular mosaicism. Finally, longitudinal analysis using scRNA-seq may provide a better understanding of the sequential changes in cellular function that occur as the RPE population matures.

## 4. Materials and Methods

### 4.1. Mice and colony management

C57BL/6J (B6) mice were produced from an animal colony bred at The Jackson Laboratory (JAX, stock #000664). Mice were housed in the Research Animal Facility at JAX on a 12-h light/12-h dark cycle and provided an NIH31 (6% fat) diet and acidified water *ad libitum*. All animal procedures were approved by the Institutional Animal Care and Use Committees of the JAX and adhered to the ARVO Statement for the Use of Animals in Ophthalmic and Vision Research.

### 4.2. RPE cell isolation and scRNA-seq

Mice at P36 were sacrificed by asphyxiation with carbon dioxide and cervical dislocation. Eyes were enucleated and placed immediately in a dish of 1x phosphatebuffered saline (PBS) on wet ice. Eyes were punctured below the limbus with a 20 g needle, gripped at the cornea with straight forceps, and cut circumferentially below the limbus with angled Vannas microdissection scissors. After removing the cornea, iris, and lens, and trimming the optic nerve flush with the sclera, the retina was immediately peeled from the posterior eyecup. The peeled eyecup was then placed in 1.0 ml prewarmed 0.5% trypsin-EDTA (Thermofisher, 15400-054) and incubated for 30 min at 37 °C under an atmosphere containing 5% CO2. Following incubation, the interior of the eyecup was grasped at the optic nerve with forceps and the entire eyecup was drawn out of the solution and resubmerged repeatedly to release RPE sheets. When no additional sheets were released, the remaining tissue containing the choroid and sclera was discarded. The solutions containing RPE sheets from both eyes were pooled with 8.0 ml of ice-cold collection buffer in a gentleMACS C tube (Miltenyi Biotec, Bergisch Gladbach, North Rhine-Westphalia, Germany) and disrupted using protocol C of a gentleMACS Dissociator (Miltenyi Biotec). Samples were centrifuged at 300*g* for 10 min at 4 °C and resuspended by triturating the pellet in 250 µl DMEM (Thermo Fisher Scientific, 11054-020) containing 2% fetal bovine serum, 2 mM EDTA. Calcium-AM (0.63 µl) was added from a 40 µM working stock and samples were incubated for 15 min. Cells negative for DAPI and positive for calcein-AM staining were sorted into DMEM contain 20% fetal bovine serum, 1x N1 medium supplement (Sigma N6530-5BL), 1x MEM non-essential amino acids (Thermo Fisher Scientific, 11140050), 2 mM GlutaMAX-I (Thermo Fisher Scientific, A128601), 0.25 mg/ml taurine (Millipore Sigma T0625), 20 ng/ml hydrocortisone (Millipore Sigma, H0396), and 13 ng/ml triiodothyronine (Millipore Sigma, T5516) using a 100 µm nozzle on a FACSAria II cell sorter (BD Biosciences, San Jose, CA, USA). Collected samples were centrifuged at 300*g* for 10 min at 4 °C and resuspended in 0.04% bovine serum albumin. A volume of the suspension containing 8,000–12,000 cells was loaded in a single channel of a Chromium Single Cell Instrument (10x Genomics, Pleasanton, CA, USA) and bar-coded cDNA libraries were prepared using a Chromium Single Cell 3’ Chip Kit v2 (10x Genomics) according to the manufacturer’s protocol. Amplified cDNA from each channel was used to construct a sequencing library (Illumina, San Diego, CA, USA). cDNA and libraries were checked for quality on a 4200 TapeStation (Agilent Technologies, Santa Clara, CA, USA) and quantified by KAPA qPCR. Sequencing was performed on an NextSeq500 System (Illumina, San Diego, CA, USA) using 150-cycle sequencing to an average depth of 50,000 reads per cell.

### 4.3. Single-cell RNA sequencing analysis

Single-cell RNA-seq data were processed using Cell Ranger v1.2.0 (10x Genomics; RRID:SCR_017344). Sequencing libraries were demultiplexed to individual cell FASTQ files utilizing cellranger mkfastq function. Each library was aligned to an indexed GRCm38 (mouse) RefSeq genome with default parameters, followed by barcode counting, and UMI counting using Cell Ranger. In R1, 2,844 single cells were sequenced with ∼191,744 mean reads and a median of ∼2,111 detected genes per cell. In R2, 2,944 single cells were sequenced with ∼128,744 mean reads and a median of ∼1,813 detected genes per cell.

Downstream analysis was performed on filtered feature counts generated by Cell Ranger. SoupX v1.5.2 [82] was implemented to estimate and remove ambient mRNA contamination. We identified potential single-cell doublets using DoubletFinder v2.0.3 [83], with an expectation of a 4% doublet rate assuming Poisson statistics, as per the developer’s code on GitHub. Further low-quality single cells containing <500 expressed genes or >5% mitochondrial transcripts were excluded from the analysis. Additionally, genes expressed in fewer than three single cells were also removed. Following the removal of low-quality and doublet cells, single cells were normalized and clustered using Seurat V4.0.0 (RRID:SCR_016341) [84]. Single-cell gene expression counts were normalized following a global-scaling normalization method with a scale factor of 10,000 and log transformed using the Seurat NormalizeData function.

We applied principal component analyses to reduce the dimensionality of the data using the top 2,000 most variable genes in the dataset. The top 10 principal components selected using the JackStraw and Elbow plot method were used in the RunUMAP analysis. Resolution parameters 0.6 and 0.5 were selected using clustering tree method [40] to identify clusters from R1 and R2 single cell datasets, respectively. Seurat V4.0.0 was used to identify cluster-specific marker genes and visualization with dot and feature plots. The genes specifically expressed in each cluster were examined to identify the cell types.

Separately, we also used an R package CELL-ID [48] to identify the cell types in our datasets. CELL-ID extract unbiased per-cell gene signature in a single-cell RNA-seq dataset and match cells from same cell type across independent datasets or reference datasets. In this study we have matched top 100 gene signatures from each cells in our single-cell RNA-seq dataset with gene signatures from melanocytes cell types [49] and microglial cell types [46] from previous studies.

### 4.4. Correlation Analysis

We have computed the Pearson correlation between our mouse RPE cell clusters and with mouse retinal cell clusters [46] using cor.test function in R. Correlations that were significant at nominal p < 0.05 are exhibited in correlation plots.

### 4.5 Functional enrichment analysis

Functional enrichment analysis was performed using the R package clusterpro-filer [85]. Gene ontology analysis was performed using enrichGO functions from the clusterprofiler R package. The function compareCluster from this package was used to compare enriched functional categories of each gene cluster. The significance threshold for all enrichment analyses was set to 0.05 using Benjamini-Hochberg corrected p-values.

### 4.6. Gene expression clustering of Marker Genes

Gene expression clustering of differentially expressed genes from each RPE cell cluster were performed using K-means clustering approach. We have used relative expression (log2FC) of differentially expressed genes in each cluster to group gene with similar expression profile across RPE cell clusters.

## Supplementary Materials

The following supporting information is available: Supplementary Table 1: Human and mouse transcriptome studies; Supplementary Figure 1: Correlation between RPE clusters; Supplementary Figure 2: Heterogeneity of RPE cell populations from R2; Supplementary Figure 3: Characterization of RPE subpopulation from R2; Supplementary File 1: Marker genes of C1-6; Supplementary File 2: GO analysis of C1-6; Supplementary File 3: k-means clustering; Supplementary File 4: GO analysis of k-means clustering groups; Supplementary File 5: Average RNA count of individual clusters and combined clusters C2–6.

## Author Contributions

Conceptualization, J.K.N., P.R., P.M.N., and G.W.C.; methodology, R.P., M.P.K., M.T.B., J.R.C., P.R.; software, R.P. and M.T.B.; validation, R.P.; formal analysis, R.P. and M.T.B.; investigation, M.T.B., J.R.C., and M.P.K.; resources, P.R. and P.M.N.; data curation, R.P. and M.T.B.; writing—R.P. and M.P.K.; writing—review and editing, M.P.K., J.K.N., P.R., P.M.N., and G.W.C.; visualization, R.P., M.T.B., and M.P.K.; supervision, J.K.N., P.R., P.M.N., and G.W.C.; project administration, P.M.N.; funding acquisition, J.K.N., P.R., P.M.N., and G.W.C. All authors have read and agreed to the published version of the manuscript.

### Funding

Research in this publication was supported by the National Eye Institute of the National Institutes of Health under award numbers R01EY011996 to P.M.N., R01EY027305 to P. M. N. and M.P.K., R01EY027860 to P. M. N. and G. W. C., and R01EY028561 to J. K. N., and by The Jackson Laboratory, Director’s Innovation Fund (DIF), award number 19000-16-13. The authors also wish to acknowledge the support of the JAX Cancer Center Single Cell Biology service, supported by the National Cancer Institute of the National Institutes of Health under award number P30CA034196.The funders had no role in study design, data collection and analysis, decision to publish, or preparation of the manuscript.

## Institutional Review Board Statement

The animal study protocol was approved by the JAX Institutional Animal Care and Use Committee (protocol ACUC 99089, last approved 22 August 2020).

## Data Availability Statement

Data are contained within the article or supplementary material. The scRNA-seq datasets will be made available through the Gene Expression Omnibus (GEO).

## Acknowledgments

The authors wish to acknowledge the support of JAX Single Cell Biology service.

