## Supplementary Figures 1-3 for "Single-cell RNA sequencing reveals molecular features of postnatal maturation in the murine retinal pigment epithelium"

**Supplementary Figure 1: Correlation between RPE clusters. (A)** Pearson correlation between RPE cell clusters from R1 mice using the average expression of genes in each cluster **(B)** Pearson correlation between RPE cell clusters from R1 mice using differential expression (log_2_FC) of genes in each cluster relative to cells in other clusters. **(C)** Pearson correlation between RPE cell clusters from R2 mice using average expression of genes in each cluster **(D)** Pearson correlation between RPE cell clusters from R2 mice using differential expression of genes in each cluster relative to cells in other clusters. Positive correlations are shown in red and negative correlations in blue color. Correlation with nominal p-value < 0.05 are considered significant.

**Supplementary Figure 2: Heterogeneity of RPE cell populations from R2 mice. (A)** Top 20 differentially expressed genes in clusters, ranked by FDR, are shown in the heatmap. Gene expression values were centered, scaled, and transformed to a scale from −2 to 2. Select signature genes are highlighted on the right. **(B)** Correlation between single cell clusters from R2 and microglial retinal cell clusters**.** Pearson correlation coefficients were calculated log fold change in expression of genes in each cluster**.** Positive correlations are shown in red and negative correlations in blue color. Correlation with nominal p-value < 0.05 are considered significant and shown in figure. **(C)** Dot plot showing marker gene expression for different RPE specific pathways (visual cycle, melanogenesis), and cell types (stem cell and immune cells). Dot sizes indicate the percentage of cells in each cluster expressing the gene, and colors indicate average expression levels. **(D)** Differential expression (log_2_FC) of melanogenesis genes along RPE clusters C1–6. **(E)** Differential expression (log_2_FC) of visual cycle genes in C1–6.

**Supplementary Figure 3: Characterization of RPE subpopulation from R2 mice. (A)** k-means clustering of differentially expressed genes in RPE cell populations from R2 mice. Clustering analysis identified 10 group of genes with distinct expression profiles across possible maturation timeline, each plotted in a different color. Number of genes in each group are shown in parentheses **(B)** Enrichment of biological processes in selected groups using clusterprofiler. The significance threshold for all enrichment analyses was set to 0.05 using Benjamini-Hochberg corrected p-values.

**Supplementary Figure 1**

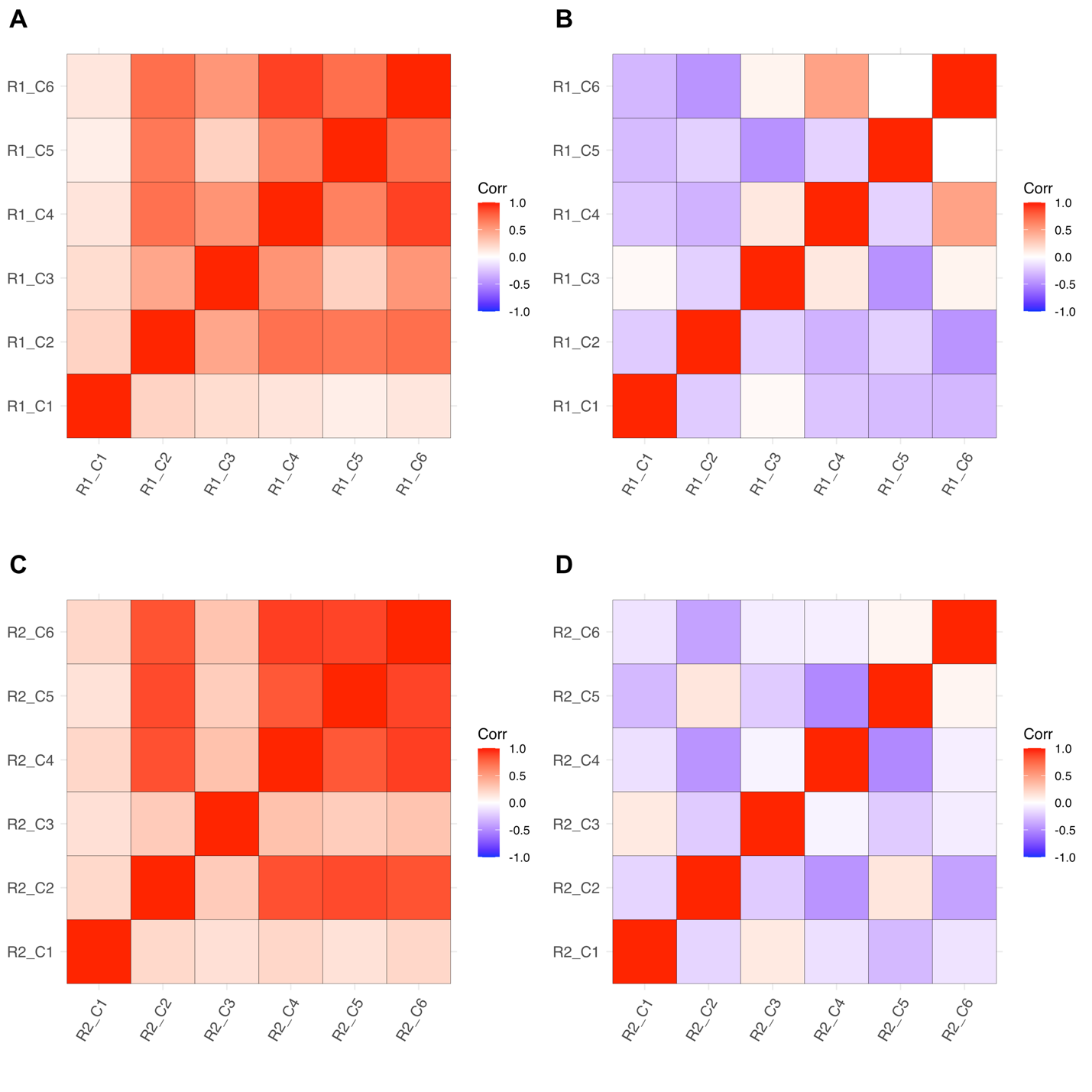

**Supplementary Figure 2**

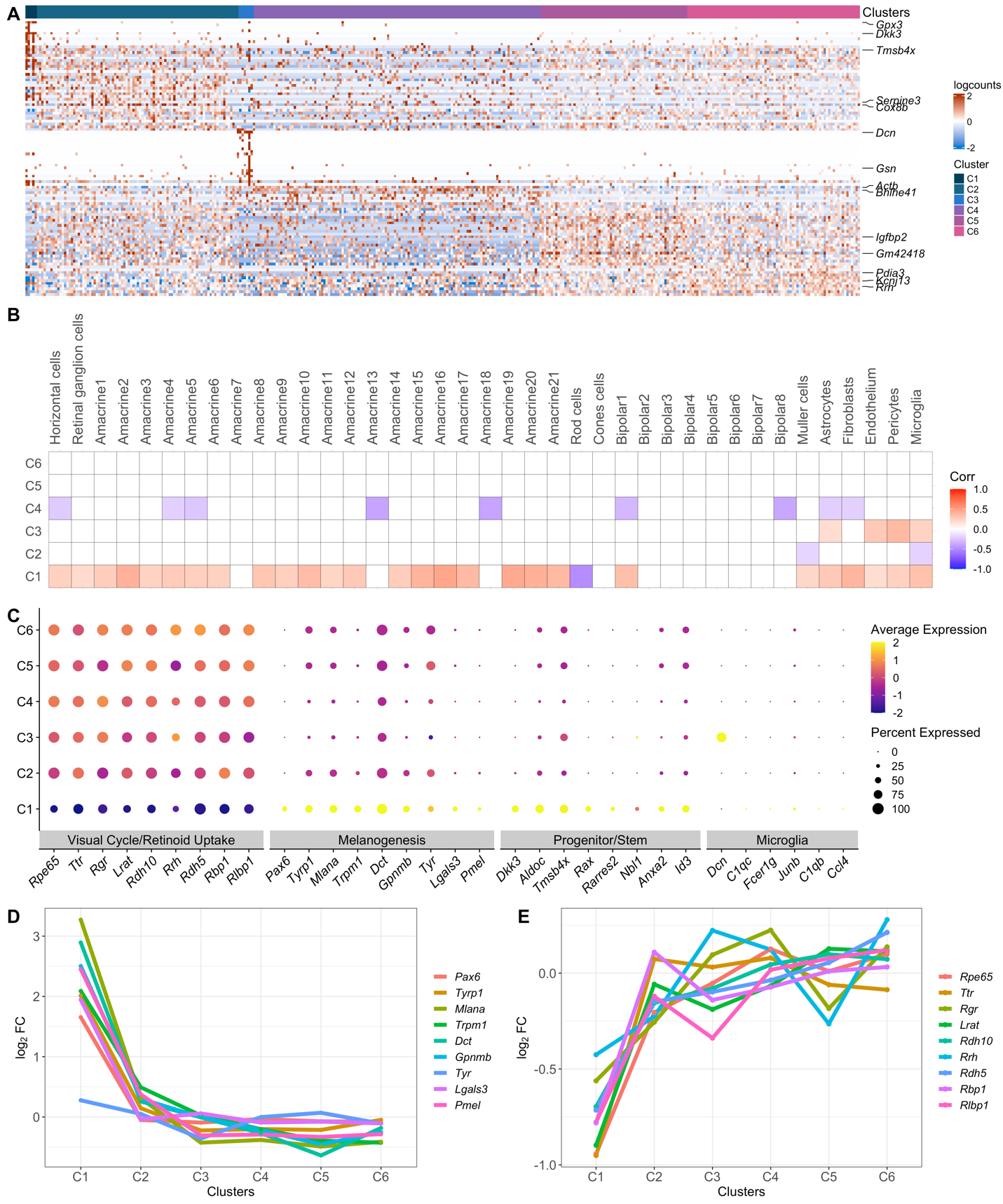

**Supplementary Figure 3**

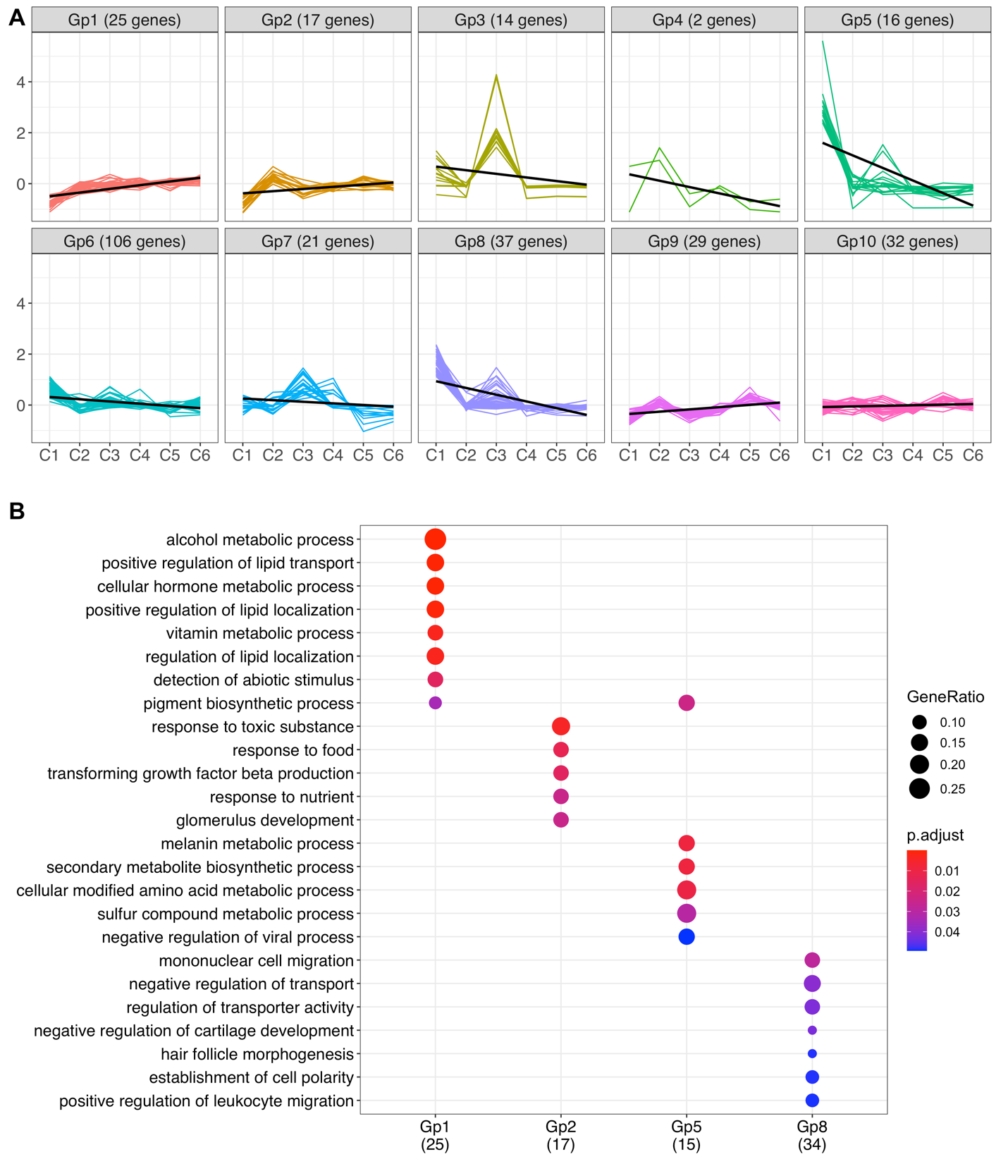
