## Supplementary Table 1 for "Single-cell RNA sequencing reveals molecular features of postnatal maturation in the murine retinal pigment epithelium"

| **Human RPE Transcriptome Studies** | | | |
| --- | --- | --- | --- |
| **Samples** | **Platform** | **Results** | **Reference** |
| 17 human eyes ages 28–66 <6h to processing, stored in RNA*later* | SAGE | Three differential expression lists were reported: macula retina enriched, periphery retina enriched, and RPE enriched. | [1] |
| 15 human eyes normal eyes (ages 45–65 and 75–87) ○ 9 males, 1 female AMD eyes (ages 78–92) ○ 3 males, 2 females from each eye, one macular and one peripheral punch of the RPE/choroid complex was taken 0.5 to 7.4 h to processing, stored in RNA*later*, stored up to a week | Microarray | 76 genes differentially expressed between macular and peripheral regions. 29 genes selected for validation via qPCR and 21 confirmed. | [2] |
| 6 human eyes ages 17–36 2 females, 4 males from each eye, one macular and two peripheral fragments were isolated 24+ h to processing | Microarray | 438 genes differentially expressed between macular and peripheral regions. Validation of 33 genes using RT- PCR resulted in an 84% correlation. | [3] |
| 6 human eyes ages 63–78 macular RPE isolated 30+ h to processing | Microarray | Attempted to identify the expression profile of macular RPE and demonstrated the lack of knowledge regarding the RPE transcriptome. Functional analysis revealed enrichment for pathways such as oxidative phosphorylation and ATP synthesis. | [4] |
| 5 human eyes ages 63–78 isolated RPE, choroid, and photoreceptor layers 16–22 h then frozen | Microarray | Identified 114 RPE specific genes. Selected 39 for validation. 85% validated through literature and s-QPCR confirmed RPE expression for remaining genes. | [5] |
| 4 adult Caucasian (ages 64–89) 4 fetal native RPE samples 4 fetal native choroid samples <24 h to processing fetal RPE primary cultures ARPE-19 cell line | Microarray | 154 RPE signature genes identified. Using qRT-PCR, 48 genes were highly expressed both in vivo and in vitro. Demonstrated that culturing cells can change the expression but did not affect enrichment of signature genes. | [6] |
| 8 normal human  RPE/choroid/sclera <6 h to processing, stored in *RNAlater*, shipped overnight, processed upon arrival | RNA-Seq | 926 genes were differentially expressed between macular and peripheral RPE/choroid/sclera | [7] |
| 4 normal human RPE/choroids <6 h to processing, stored in liquid nitrogen nasal, temporal, and macular regions | RNA-Seq | 81 genes showed increased expression in the nasal RPE/choroid and 39 genes showed decreased expression | [8] |
| RPE-choroid from 6 donor eyes | scRNA-Seq |  | [9] |
| RPE from 3 donor eyes | scRNA-Seq |  | [10] |

| **Mouse RPE Transcriptome Studies** | | | |
| --- | --- | --- | --- |
| **Samples** | **Platform** | **Results** | **Reference** |
| Compare C57BL/6J mouse and human RPE versus photoreceptors and choroid isolated by laser microdissection. Look for signature genes, correcting for possible RNA contamination from its adjacent layers. | Microarray |  | [11] |
| Mice were examined at 6 time points. RPE in n=5 male C57BL/6NCrl mice (10–13 weeks of age) per time point was scraped from peeled eyecups in RNA*later*. | RNAseq | Expression of genes encoding tight junction protein were found to vary with circadian rhythm (Zeitgeber time (ZT) 0, 2, 4, 9, 14, and 19). | [12] |
| Two eyes pooled from n=3 mice at two time points with varying light-dark history. Retinas were peeled, RNA*later* was added, and sonication was used to release cells. | RNAseq | 14,083 transcripts were shared between the cohorts. | [13] |
| Mice were examined at 6 time points. RPE in n=5 male C57BL/6NCrl mice (10–13 weeks of age) per time point was scraped from peeled eyecups in RNA*later* . | RNAseq | Found 756 significant DEGs in murine RPE when comparing six different ZT points. The highest energy demand of RPE cells is at night, whereas POS phagocytosis and degradation take place in the morning. Fatty acid and glycerophospholipid synthesis genes are upregulated at night, possibly playing a role in generating building blocks for membrane synthesis. | [14] |
